# Characterisation of the bovine C-type lectin receptor Mincle and potential evidence for an endogenous ligand^1^

**DOI:** 10.1101/2023.03.20.533273

**Authors:** Angela Holder, Jeannine Kolakowski, Chloe Rosentreter, Ellen Knuepfer, Sabine A. F. Jégouzo, Oliver Rosenwasser, Heather Harris, Lotta Baumgaertel, Amanda Gibson, Dirk Werling

**Author notes:** Correspondence: Prof. Dirk Werling. Contributed equally to this work (Angela Holder, Jeannine Kolakowski).

## Abstract

Innate immune receptors that form complexes with secondary receptors, activating multiple signalling pathways, modulate cellular activation and play essential roles in regulating homeostasis and immunity. We have previously identified a variety of bovine C-type lectin-like receptors that possess similar functionality than their human orthologues. Mincle (*CLEC4E*), a heavily glycosylated monomer, is involved in the recognition of the mycobacterial component Cord factor (trehalose 6,6′-dimycolate). Here we characterise the bovine homologue of Mincle (boMincle), and demonstrate that the receptor is structurally and functionally similar to the human orthologue (huMincle), although there are some notable differences. In the absence of cross-reacting antibodies, boMincle-specific antibodies were created and used to demonstrate that, like the human receptor, boMincle is predominantly expressed by myeloid cells. BoMincle surface expression increases during the maturation of monocytes to macrophages. However, boMincle mRNA transcripts were also detected in granulocytes, B cells, and T cells. Finally, we show that boMincle binds to isolated bovine CD4^+^ T cells in a specific manner, indicating the potential to recognize endogenous ligands. This suggests that the receptor might also play a role in homeostasis in cattle.

## Introduction

C-type lectin receptors (CLRs) are pattern recognition receptors that play a dual function: they regulate immune cell homeostasis and sense glycans expressed by pathogens to induce innate immune responses such as phagocytosis, antigen presentation, cytokine production, and subsequent T cell activation (1).

The “macrophage inducible C-type lectin” (Mincle), encoded by the *CLEC4E* gene, is a member of the Dectin-2 family of CLRs, which includes Dectin-2, blood dendritic cell antigen 2 (BDCA-2, CD303), dendritic cell immunoreceptor (DCIR), dendritic cell immunoactivating receptor (DCAR), and macrophage C-type lectin (MCL). Like the other members of this family, Mincle is a type II transmembrane protein with a single extracellular carbohydrate recognition domain (CRD) (2). Following ligand binding to the CRD, the cellular responses induced by CLRs are generally either activating or inhibitory, depending on the presence of an immunoreceptor tyrosine-based activation motif (ITAM) or an immunoreceptor tyrosine-based inhibitory motif (ITIM) in their cytoplasmic domain (3). However, some CLRs, including several members of the Dectin-2 family, do not possess either motif and must associate with a signalling adaptor molecule (1). Mincle uses a conserved arginine residue (Arg42) in its transmembrane domain to associate with the ITAM-bearing FcRγ chain (4). Phosphorylation of the FcRγ-linked ITAM motif upon ligand binding to Mincle recruits SH2 domain-containing protein tyrosine kinases (e.g., Syk). Syk can activate the Caspase recruitment domain family member 9 (Card9) – B-cell CLL/lymphoma 10 (Bcl10) – mucosa-associated lymphoid tissue lymphoma translocation protein 1 (MALT1) signalosome which results in activation of the transcription factor NF-kB. Such upregulated transcription leads to the production of immunoregulatory cytokines and chemokines including IL-6, IL-12, IL-23, and IL-1ß (5).

Studies in humans and mice have demonstrated that Mincle is expressed on the surface of monocytes (Mo), macrophages (M0), dendritic cells (DC), neutrophils, and B cells (4, 6-8), and transcribed in T cells (9). The receptor was initially identified as a molecule whose transcription can be induced by lipopolysaccharide (LPS) via the transcription factor NF-IL6 (10). Subsequently, Mincle has been found to recognise a wide variety of targets and ligands, both endogenous and exogenous (11). To maintain self-homeostasis, Mincle can recognise lipid-based damage-associated molecular patterns (DAMPs), thereby monitoring the internal environment. Exposure to some pathogenic fungi, including *Candida (C*.*) albicans* and several species of *Malassezia (M*.), has been shown to up-regulate Mincle expression on the surface of macrophages (12, 13). Furthermore, *C. albicans* infections in Mincle-deficient mice resulted in a significantly higher fungal burden compared to wild-type mice (13), while co-cultures of *M. pachydermatis* with bone marrow-derived macrophages (BMDM) from Mincle-deficient mice resulted in a significant disruption of the pro-inflammatory cytokine response (14). The ligand expressed by *Malassezia spp*. was shown to be a glyceroglycolipid (15). In *Mycobacterium spp*. the main ligand of Mincle is trehalose-6,6’-dimycolate (TDM), a glycolipid found in the outer membranes of the bacterium (16, 17). Binding of TDM is facilitated by Ca^2+^ while binding of spliceosome-associated protein 130 (SAP130), a component of the U2 small nuclear ribonucleoprotein complex, which is released by nuclei of dead and damaged cells after translocation to the external milieu (4), is Ca^2+^-independent. Whether post-translational modifications of SAP130 confer the ability of Mincle to bind SAP130, or whether Mincle recognises SAP130 peptides is currently unclear.

Interactions of the human (hu)Mincle CRD with trehalose as well as mono- and diacylated derivatives of trehalose have been previously investigated (18). Comparisons of these interactions with those of CRDs from murine (mu) and bovine (bo)Mincle revealed similarities and species-specific differences (19, 20), which might contribute to the different responses of human and bovine M0 infected with either Mtb or *M. bovis* (21). Given these species-specific differences, the aim of the present study was to characterise the function and ligands of boMincle further.

## Materials and Methods

### Cell isolation and culture

Bovine peripheral blood mononuclear cells (boPBMCs) were purified by density gradient centrifugation (Lymphoprep, STEMCELL technologies) using whole blood from Holstein Friesian cattle in mid lactation reared at the Royal Veterinary College (Hatfield, UK) and sampled under home office licence (PPL7009059). Granulocytes were purified from the erythrocyte-containing layer of the density gradient by hypertonic lysis of the red blood cells in 20 ml distilled water followed by a return to isotonic conditions by the addition of 2 ml 10 % NaCl. After several washes in PBS (Gibco), the cells were placed in tissue culture medium (TCM) consisting of RPMI-1640 medium with Glutamax-1 (Gibco), supplemented with 10 % foetal bovine serum (Sigma-Aldrich) and 1 % Penicillin-Streptomycin (Gibco).

Non-cultivated, freshly isolated boPBMCs were fluorescently labelled for subsequent separation of T cells, B cells, and Mo by fluorescence-activated cell sorting (FACS). Cells were labelled with a receptor-specific primary antibody and a fluorescently labelled secondary antibody, or with directly conjugated primary antibodies directed towards cell-specific surface markers (Suppl. Tab. 1). Cell staining was performed in PBS / 1 % goat serum for 40 min at 4°C after blocking in PBS / 3 % goat serum for 20 min at 4°C. Specificity controls included unstained cells and PBMCs labelled with the secondary antibody alone. Stained PBMCs were resuspended in 500 µl FACSFlow buffer (BD Biosciences) and sorted on a FACSAria Fusion (BD Biosciences). The machine was equipped with four lasers (355nm, 405nm, 488nm, 640nm) and standard filters (EP: 525/50, 450/50, 488/10, 530/30, 585/42, 610/20, 616/23, 660/20, 670/30, 695/40, 710/50, 730/45, 780/60). Acquisition settings were determined using the specificity controls, at least 10’000 events were recorded in dot plots using the BD FACSDiva V9 software (BD Biosciences), and data were analysed using FlowJo V10 (BD Biosciences). Bovine T cells, B cells, and Mo were sorted separately into 15 ml Falcon tubes. An example of the gating strategy is shown in (Suppl. Fig. 1). Separated bovine leukocyte subsets were washed in PBS, lysed in RIPA buffer (Sigma-Aldrich), and stored at -80°C for RNA extraction.

Monocyte-derived macrophages (MDMØ) were generated from boPBMCs plated into six-well plates at a concentration of 6 × 10^6^ cells / well in 3 ml TCM. Cells were incubated at 37°C and 5 % CO_2_ for 1-2 h until the Mo had adhered to the plastic. Plates were washed twice with PBS to remove any non-adherent cells before the addition of TCM containing 20 ng mL^-1^ recombinant bovine (rbo)M-CSF (Kingfisher Biotech). Every 2-3 days 2 ml / well of spent media was replaced with fresh TCM containing rboM-CSF. MDMØ were harvested using Accutase (Gibco). MDMØ from human blood were generated as described above. Briefly, a total of 50 ml of whole human blood was collected by a trained phlebotomist from healthy donors in BD Vacutainer^®^ EDTA tubes (BD Vacutainer^®^, USA). This was previously approved by the RVC’s Ethics and Welfare Committee (URN 2019 1916-3).

### Comparative analysis of bovine *CLEC4E*/Mincle sequences and protein models

The chromosomal location, DNA, and amino acid sequences for bovine *CLEC4E*/Mincle were extracted from the current bovine genome annotation (ARS-UCD1.2) using the NCBI gene database (https://www.ncbi.nlm.nih.gov/). BLAST searches of the bovine genome (https://blast.ncbi.nlm.nih.gov/-Blast.cgi) using the nucleotide sequences for human and murine Mincle (NM_014358.4 and NM_019948.2) were used to confirm that only one copy of the gene existed. Sequences for sheep, goat, pig, and horse *CLEC4E*/Mincle (XM_027967911.1, XM_005680910.3, XM_021092401.1 and XM_001492794.4) were also obtained from the NCBI gene database for comparison. Sequence alignments were generated using CLC Main Workbench 22 (Qiagen). The percentages of sequence identity were calculated by the pairwise sequence alignment programme EMBOSS Needle (https://www.ebi.ac.uk/Tools/psa/emboss-_needle/).

Amino acid sequences for bovine (XP_010803863.1), murine (NP_064332.1), and human (NP_055173.1) Mincle were obtained from the NCBI database. CRD sequences for each protein were identified using the Simple Modular Architecture Research Tool (SMART) (http://smart.embl-heidelberg.de). 3D protein structures for each CRD were constructed with the template-based automated modelling algorithm of SWISS-MODEL (https://swissmodel.expasy.org). Multiple CRD structure superimpositions were performed using the ChimeraX-1.2.5 software. CRD model characteristics that were compared in the analysis include the overall structure, hydrophobicity, and electrostatic surface potential.

RNA was extracted from PBMCs of 10 Holstein Friesian cattle using the RNeasy Mini Kit (Qiagen) according to the manufacturer’s instructions. Subsequent cDNA synthesis was conducted using the iScript cDNA Synthesis Kit (Bio-Rad). Gene-specific primers were designed to amplify the coding region of bovine *CLEC4E* (Mincle) (Suppl. Tab. 2). PCR reactions were performed using Phusion High-Fidelity Master Mix (Thermo Fisher Scientific). The resulting PCR products were separated by agarose gel electrophoresis, purified using the QIAquick Gel Extraction Kit (Qiagen), and submitted for Sanger DNA sequencing (MRC PPU DNA Sequencing Service, Dundee University).

### Detection of *CLEC4E/*Mincle expression in bovine leukocyte subsets by RT-PCR

RNA was extracted from cell lysates of bovine granulocytes and sorted PBMC subsets of three to seven Holstein Friesian cattle using the AllPrep RNA/Protein kit (QIAGEN) according to the manufacturer’s instructions. cDNA synthesis from extracted RNA was performed using the iScript cDNA synthesis kit (Bio-Rad) with modified reaction conditions (priming at 25°C for 5 min, 2x reverse transcription at 42°C for 30 min, RT inactivation at 85°C for 5 min). Quality control of synthesised cDNA was performed by PCR with gene-specific primers amplifying bovine β-actin (Suppl. Tab. 2). PCR reactions were performed using OneTaq Quick Load 2X Master Mix with Standard Buffer (New England Biolabs). Detection of Mincle-specific cDNA was conducted by PCR with the same gene-specific primers described in the previous section (Suppl. Tab. 2). The resulting PCR products were separated by agarose gel electrophoresis and visualised using a G-Box illuminator (Syngene).

### Synthesis of bovine Mincle CRD fragments

Bovine Mincle CRD fragments, with and without a biotin tag, were generated as described previously (19).

### Generation of recombinant anti-bovine Mincle antibodies

HuCAL anti-bovine Mincle CRD antibodies were generated commercially (BioRad HuCAL Platinum).

### Confirmation of immunogen and protein binding by anti-bovine Mincle CRD antibodies

Accurate binding specificities of custom-made, anti-bovine Mincle CRD antibodies were assessed by SDS-PAGE and Western Blot analysis. Recombinant bovine Mincle CRD (immunogen) served as positive control. Isolated bovine CD14^+^ Mo and boPBMCs excluding CD14^+^ Mo were solubilised in Laemmli sample buffer with or without DTT and separated on 12 % Bis-Tris Gels (Thermo Fisher Scientific) using MES (immunogen) (Thermo Fisher Scientific) or MOPS (boPBMCs) (Thermo Fisher Scientific) SDS running buffer. Proteins were transferred onto nitrocellulose membranes (Thermo Fisher Scientific) using NuPage transfer buffer (Thermo Fisher Scientific), and membranes were blocked using skimmed milk (5 %) in PBS with 0.1 % Tween20 overnight. Blocked membranes were incubated with HuCAL anti-bovine Mincle CRD antibodies (2.5 µg mL^-1^) before incubation with secondary goat-anti-human IgG F(ab’)2: HRP (STAR126P, Bio-Rad) antibody. Signals were detected by enhanced chemiluminescence (ECL) using Luminata Forte (Millipore). ECL signal and protein standard were detected using a G-box illuminator (Syngene).

### Expression of C-type lectin receptors in Chinese Hamster Ovary (CHO) cells

cDNA was generated from boPBMCs as mentioned above. Full-length sequences for bovine *CLEC4E* (Mincle) and *CLEC4L* (CD209, DC-SIGN) genes were amplified by PCR with gene-specific primers (Suppl. Tab. 2) using Phusion High-Fidelity Master Mix (ThermoFisher). PCR products were digested with AgeI and NheI, purified by gel extraction (QIAquick Gel Extraction Kit, Qiagen), and ligated into pUNO1 plasmid (Invivogen) using T4 DNA ligase (Thermo Fisher Scientific). Successful cloning was confirmed by Sanger DNA sequencing (MRC PPU DNA Sequencing Service, Dundee University). 1 × 10^5^ CHO cells were seeded per well in a 24-well plate in supplemented Minimal Essential Medium (MEM with 10 % FBS, 2 % non-essential amino acid solution (Sigma Aldrich), 1 % GlutaMAX-1, 1 % Penicillin-Streptomycin (Gibco)), and cultured overnight until they reached 70-90 % confluency. These confluent CHO cells were transfected with 700 ng pUNO1-Mincle or pUNO1-DC-SIGN using TurboFect Transfection Reagent (Thermo Fisher Scientific) according to the manufacturer’s instructions. C-type lectin receptor expression was confirmed by flow cytometry.

### Analysis of Mincle surface expression on CHO cells, PBMCs, MDMØ by flow cytometry

Cells were labelled indirectly with receptor-specific primary antibodies and a fluorescently labelled secondary antibody (Suppl. Tab. 1), and/or with a directly conjugated antibody directed towards a cell surface marker (Suppl. Tab. 1). Staining was conducted in PBS / 1 % BSA (CHO cells and huPBMCs) or PBS / 1 % BSA / 1 % normal mouse serum (bovine cells) (Sigma-Aldrich) for 40 min at 4°C. Bovine cells were blocked ahead of antibody staining in PBS / 1 % BSA / 5 % mouse serum for 20 min at 4°C. Specificity controls included unstained cells as well as cells labelled with isotype controls or the secondary antibody alone (Suppl. Tab. 1). Viability of PBMCs was controlled by staining with Annexin V CF-Blue (Abcam). For CHO cells and MDMØ, trypan blue exclusion was performed showing that less than 10% cells were dead. Zymosan A (*S. cerevisiae*) bioparticles conjugated with Alexa Fluor 488 (Thermo Fisher Scientific) were used to stain CHO cells transfected with pUNO1-boDC-SIGN. Flow cytometric analysis was carried out using a LSRFortessa X-20 (BD Biosciences) equipped with four lasers (355nm, 405nm, 488nm, 640nm) and standard filters (EP: 379/28, 450/50, 488/10, 525/50, 530/30, 575/25, 610/20, 670/30, 710/50, 730/45, 780/60). Acquisition settings were determined using the specificity controls, counting a minimum of 10,000 gated cells that were recorded with the BD FACSDiva V9 software (BD Biosciences). Data were analysed using FlowJo V10 software (BD Biosciences).

### Assessment of cytokine responses in bovine MDMØ stimulated with Mincle ligands

Trehalose-6,6-dibehenate (TDB, Invivogen) was resuspended in isopropanol at 1 mg mL^-1^, heated at 60°C for 2 min, and then vortexed until dissolved. The TDB was used plate-bound by coating wells of a 48-well plate with 25 µg TDB. Mock treated wells were used as control. Bovine MDMØ were plated into the 48-well plates at a density of 1.5 × 10^5^ cells / well. Lipopolysaccharide (LPS) from *E. coli* O127:B8 (Sigma Aldrich) was resuspended at 1 mg mL^-1^ in dH_2_O and used at 1000 ng mL^-1^ (500 ng / well). HuCAL anti-bovine Mincle CRD antibodies (AbD31621 and AbD31662) and HuCAL Fab-dHLX-FH Negative control (Bio-Rad) were added at 2.5 µg mL^-1^ (1.25 µg / well). Bovine MDMØ were incubated with ligands and/or antibodies for 24 h at 37°C and 5 % CO_2_. Cell culture supernatants were then collected and stored at -20°C. Secretion of TNF-α was measured by sandwich ELISA (R&D Systems).

### *M. bovis* Bacillus Calmette-Guerin preparation

*M. bovis* Bacillus Calmette-Guerin (BCG) (Pasteur strain) expressing green fluorescent protein (GFP), kindly provided by Dr. Amanda Gibson (Aberystwyth University), was cultured at 37°C for 14 days in Middlebrook 7H9 medium (Difco, BD) containing 10 % Middlebrook ADC supplement (Difco, BD), 0.05 % Tween80 (Sigma-Aldrich), 2 % glycerol and 20 µg mL^-1^ kanamycin. Aliquots were stored at -20°C until further use. Upon defrosting, the BCG was centrifuged at 400 x g for 10 min and the pellet resuspended in PBS by vortexing. Colony forming units (CFU) were estimated as described from OD_600_ measurements (OD_600_ 1.0 = 3.13 × 10^7^ CFU mL^-1^) and used for subsequent calculation of the MOI (22).

### Phagocytosis assay

2 × 10^5^ bovine MDMØ were seeded in each well of a 96-well U bottom plate in 100 µl TCM, infected with GFP-BCG (MOI of 5) in PBS at 37°C and 5 % CO_2_ for 90 min. Cells incubated at 4°C served as control since phagocytosis was prevented under these conditions. Infection of MDMØ was followed by two washing with ice-cold PBS and killing of surface-bound bacteria by incubation with 50 µg mL^-1^ gentamicin (Sigma-Aldrich) for 30 min. After two washes cells were fixed with 4 % paraformaldehyde (Cytofix, BD) for 30 min at 4°C and analysed by flow cytometry. The role of Mincle was investigated by incubating the cells with custom-made HuCAL anti-bovine Mincle CRD antibodies (AbD31621 and AbD31662) and HuCAL Fab-dHLX-FH Negative control (Bio-Rad) at 20 µg mL^-1^ for 15 min prior to addition of GFP-BCG.

### CD4^+^ T cell binding assay

CHO cells transfected with empty pUNO1, pUNO1-boDC-SIGN, and pUNO1-boMincle were transferred to 4-well chamber slides (Lab Tek-II, Nunc) at a density of 1 × 10^5^ cells / chamber, and incubated at 37°C and 5 % CO_2_ overnight. Bovine CD4^+^ T cells were isolated from PBMCs by magnetic-activated cell sorting (MACS) using a mouse anti-bovine CD4 antibody (clone CC8, Bio-Rad), anti-mouse IgG MicroBeads (Miltenyi Biotec), and LS columns (Miltenyi Biotec), according to the manufacturer’s instructions. Successfully isolated CD4^+^ T cells were fluorescently labelled with 0.1 mM carboxyfluorescein succinimidyl ester (CFSE) and added to the chamber slides at a density of 5 × 10^5^ T cells / chamber. Transfected CHO cells and CFSE-labelled CD4^+^ T cells were incubated together at 37°C and 5 % CO_2_ for 30 min. Slides were washed in PBS, and cells visualised with a Nikon Eclipse Ti2 microscope. Image Pro Plus 5.0 software (Media Cybernetics) was used to count the number of fluorescent CD4^+^ T cells in each field of view; 8 randomly selected fields were counted per chamber. The CHO cells transfected with empty pUNO1 and pUNO1-boDC-SIGN represented negative and positive controls for T cell binding, respectively.

## Results

### Comparative analysis of *CLEC4E/*Mincle sequences

Analysis of the current bovine genome annotation (ARS-UCD1.2) identified the sequence for bovine Mincle (*CLEC4E*) as a single copy gene located in a region on chromosome 5. This region also contains numerous NK cell receptor genes, and the Mincle gene clustered with other members of the Dectin-2 family. The same has been observed for both human and mouse *CLEC4E* on the orthologous chromosomes 12 and 6, respectively (Fig. 1). Comparison of amino acid sequences of bovine *CLEC4E*/Mincle with those of human, mouse, and other livestock species (sheep, goat, pig, and horse) showed a high degree of sequence similarity (Tab. 1). As expected, comparison of bovine *CLEC4E*/Mincle sequences with those of other ruminant species such as sheep and goat yielded the highest percentage of sequence identity.

**Fig. 1.**
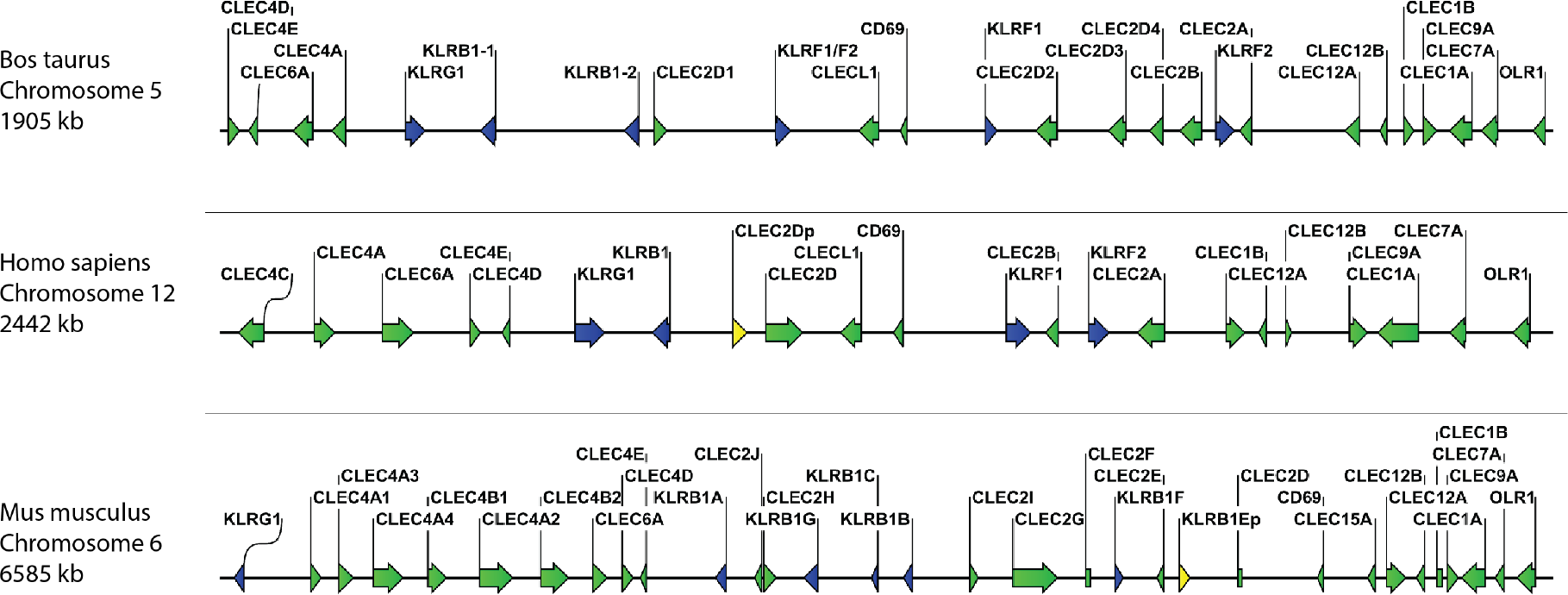
Comparative organisation of genomic regions containing C-type lectin receptor genes in selected species The genes for bovine, human, and murine Mincle (CLEC4E) as well as other Dectin-2 family CTLR (CLEC4A, CLEC4C, CLEC4D, and CLEC6A) cluster together in regions with other CTLR and NK cell receptor genes (KLR genes). However, the spacing of genes on the chromosomes varies between species. The gene for BDCA-2 (CLEC4C) is only present in humans, while mice possess multiple paralogues for the DCIR (CLEC4A) gene. CTLR genes are shown in green, killer cell lectin-like receptors (KLR) in blue, and pseudogenes in yellow. Genomic regions are not drawn to scale.

**Table 1:** Percentage of sequence identity with bovine Mincle

| Species | Identity (%) |
| --- | --- |
| Human | 71.7 |
| Mouse | 69.3 |
| Sheep | 92.0 |
| Goat | 91.0 |
| Pig | 79.7 |
| Horse | 76.4 |

### Analysis of polymorphisms in the bovine *CLEC4E* gene

Sequencing of the *CLEC4E* coding region in cDNA samples from ten Holstein Friesian cattle identified three SNPs (Tab. 2 and Suppl. Tab. 3). Two of these SNPs (rs135842138 and rs137802077) were synonymous, while the third (rs135158086) resulted in an isoleucine to threonine amino acid substitution (I174T). For all three SNPs the frequency of the alternative allele was higher than that of the reference allele, which potentially represents a general difference between beef and dairy cattle since the reference genome is from a Hereford cow.

**Table 2:**
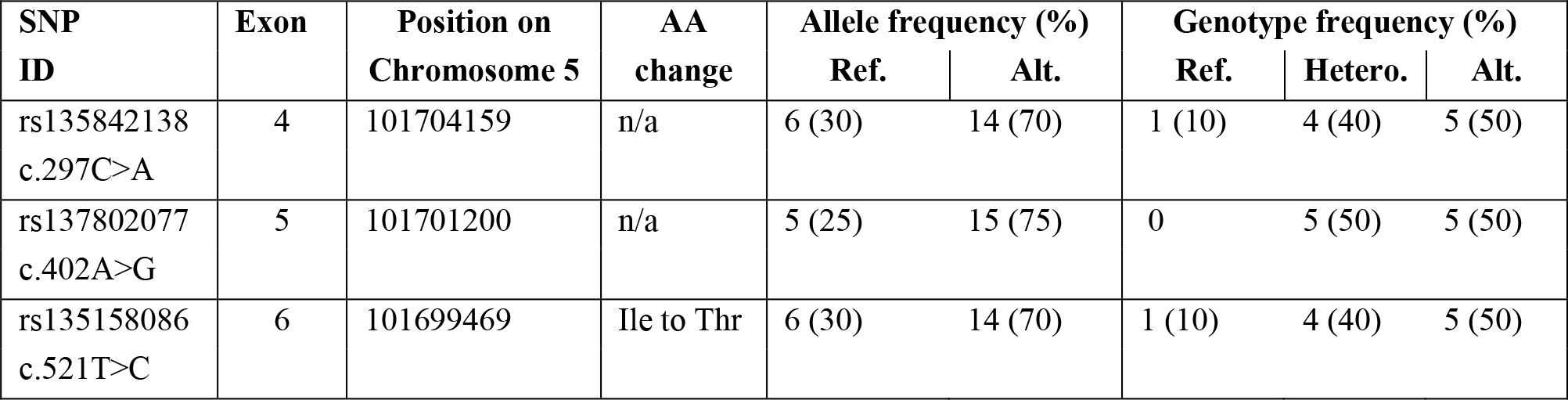
Analysis of CLEC4E coding region SNPs in Holstein Friesian Cattle. The corresponding exon and allele frequency were identified for each SNP. This information was used subsequently to estimate genotype frequencies for homo- and heterocygocyty.

| SNP ID | Exon | Position on Chromosome 5 | AA change | Allele frequency (%) |  | Genotype frequency (%) |  |  |
| --- | --- | --- | --- | --- | --- | --- | --- | --- |
|  |  |  |  | Ref. | Alt. | Ref. | Hetero. | Alt. |
| rs135842138<br>c.297C>A | 4 | 101704159 | n/a | 6 (30) | 14 (70) | 1 (10) | 4 (40) | 5 (50) |
| rs137802077<br>c.402A>G | 5 | 101701200 | n/a | 5 (25) | 15 (75) | 0 | 5 (50) | 5 (50) |
| rs135158086<br>c.521T>C | 6 | 101699469 | Ile to Thr | 6 (30) | 14 (70) | 1 (10) | 4 (40) | 5 (50) |

One of the cattle was found to possess two versions of the *CLEC4E* gene, the full-length version and a splice variant missing a 32bp region at the end of exon 5 (Suppl. Fig. 2).

### Comparative analysis of Mincle CRD protein models

A comparison of 3D protein structures of human, bovine, and murine Mincle CRDs was included to highlight structural similarities and differences of these receptor CRDs (Fig. 2). The models were generated using SWISS-MODEL’s template-based automated modelling algorithm and analysed with the ChimeraX-1.2.5 software. 3D models of the same protein but bound to different ligands (here citrate and trehalose) underline how at least part of the Mincle CRD has different conformations under different conditions (19, 23). Of particular interest is the loop with amino acid residues (Leu-172 and Val-173 in the bovine model) forming one side of the hydrophic groove that was first identified in the trehalose-bound conformation of bovine Mincle CRD (19). This groove although not present in the unbound or citrate-bound conformation of human (Fig. 2) and murine Mincle CRDs is predicted to be formed in both proteins at maximum Ca^2+^ saturation of the CRD at the right pH and binding of trehalose or acylated trehalose (18, 20). The binding mechanism of trehalose and mono-or diacylated trehalose derivatives is therefore likely to be similar between all three protein CRDs (18-20). However, structural differences between Mincle CRDs that also include the exact amino acids present in and along the hydrophobic groove adjacent to the primary carbohydrate binding site (Fig. 2, top row, orange amino acids) might still result in differential binding of other ligands. In this regard, glycerol monomycolate was identified as a ligand for human but not for murine Mincle (24).

**Fig. 2.**
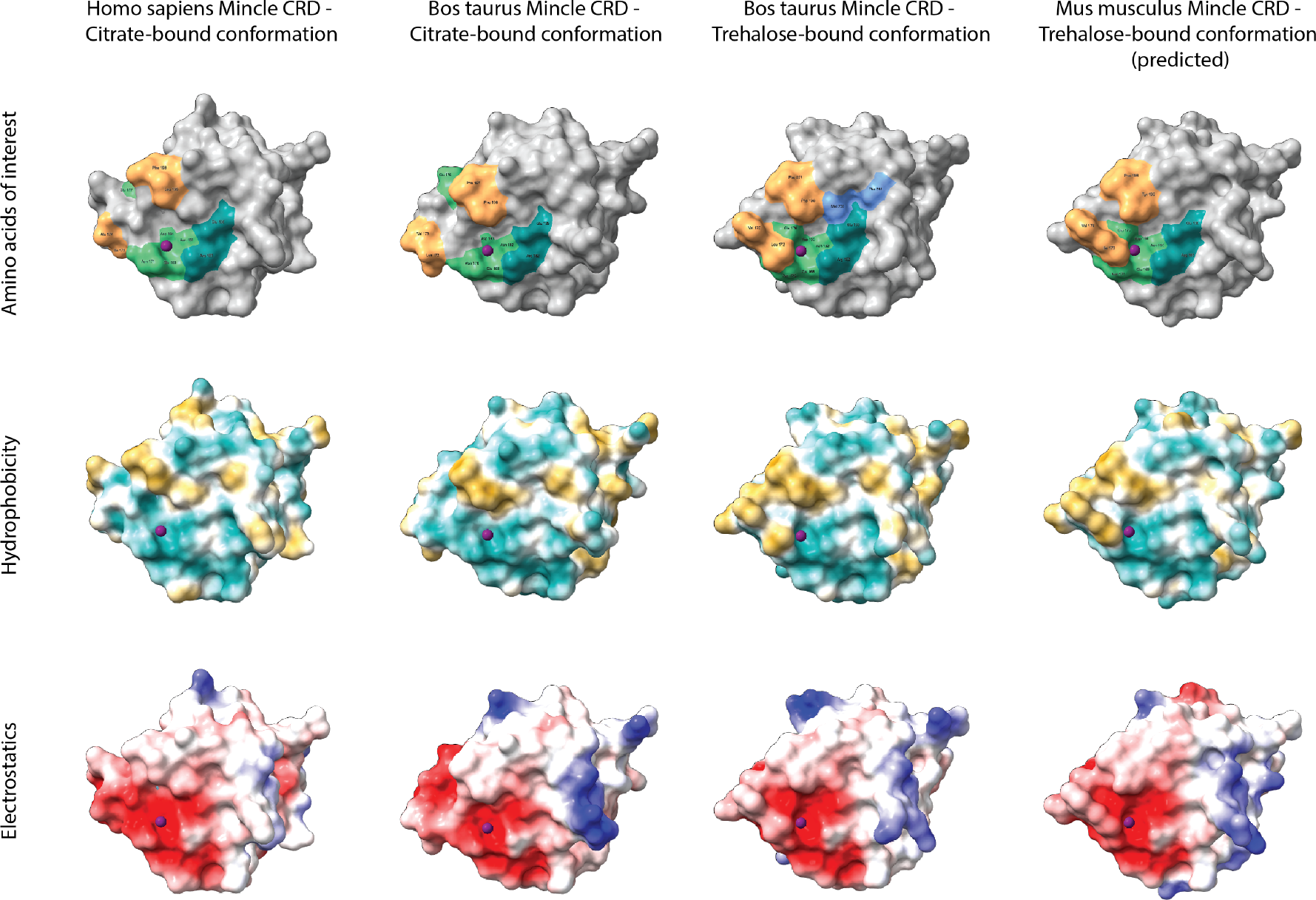
Comparative analysis of Mincle CRD protein models Tertiary structures of human (template 3wh2.1.A), bovine (templates 4kzw.2.A (citrate-bound) and 4zrw.1.A (trehalose-bound)), and murine (template 4zrw.1.A) Mincle CRDs were generated using SWISS-MODEL’s template-based automated modelling algorithm and analysed with the ChimeraX-1.2.5 software. Protein structures bound to different ligands (here citrate and trehalose) were included to underline the conformational changes of the same protein under different conditions. The top row highlights amino acid residues of each CRD that are involved in ligand binding. Light green amino acids around the purple Ca^2+^ ion form the primary carbohydrate binding site which binds glucose 1 of trehalose among other ligands. Amino acid Glu-176 (bovine models) or Glu-177 (human and murine models) thereby relocates towards or away from the primary binding site based on the interacting ligand (citrate- and trehalose-bound structural comparison). Dark green amino acids establish 2-OH group bonds with glucose 2 of trehalose and form a secondary carbohydrate binding site. Orange amino acids frame the hydrophobic groove present in the trehalose-bound conformation of bovine Mincle and likely present in the trehalose-bound conformation of human and murine Mincle (yet no X-ray crystal structure available for trehalose-bound human Mincle and no X-ray crystal structure available for any conformation of murine Mincle (here only predicted)). Both blue amino acids shown in the trehalose-bound conformation of bovine Mincle CRD represent part of an extended carbohydrate binding site that can facilitate binding of acyl chains outside of the hydrophobic groove (23). The middle row nicely demonstrates the hydrophobicity of the groove (gold) adjacent to the primary carbohydrate binding site in the trehalose-bound bovine and predicted murine Mincle CRD, while the bottom row showcases the electrostatic surface potential of each protein (negatively charged areas in red, positively charged areas in blue).

### Detection of *CLEC4E* transcripts in bovine leukocyte subsets by RT-PCR

The transcription pattern of *CLEC4E* in bovine leukocyte subsets was evaluated by RT-PCR in cDNA samples from freshly isolated, FACS-sorted T cells, B cells, Mo, and granulocytes of three to seven different Holstein Friesian cattle (Suppl Fig. 3). Interestingly, *CLEC4E* cDNA was generally detected in all four leukocyte subsets with variations between different cattle (Tab. 3). While the results for bovine granulocytes were most consistent, cDNA of *CLEC4E* was only found in lymphocytes and Mo of some of the animals tested. One potential explanation for this finding is an inconsistent expression of Mincle as an inducible C-type lectin receptor.

**Table 3:** Detection of CLEC4E transcripts in four different bovine leukocyte subsets by RT-PCR. RNA for cDNA synthesis was extracted from cells of three to seven different Holstein Friesian cattle.

| Receptor | Granulocytes | T cells | B cells | Monocytes |
| --- | --- | --- | --- | --- |
| Mincle | 3/3 | 2/7 | 4/7 | 6/7 |

### Anti-bovine Mincle HuCAL antibodies detect the immunogen and the full-length Mincle protein in SDS-PAGE and Western Blot analysis

Antibodies against boMincle were produced by panning a commercially available HuCAL phage-display library using a recombinant bovine (rbo)Mincle CRD as immunogen. Such panning identified 16 antibodies that demonstrated specific antigen binding by indirect ELISA during the screening process. This result was subsequently confirmed by SDS-PAGE and Western Blot analysis. All of the antibodies tested resulted in a clear band of the expected size when using the recombinant bovine Mincle CRD fragment (15 kDa; immunogen) (Suppl. Fig. 4), and a corresponding yet varyingly strong band for the full-length protein was identified in bovine CD14^+^ Mo (19 kDa; Fig. 3).

**Fig. 3.**
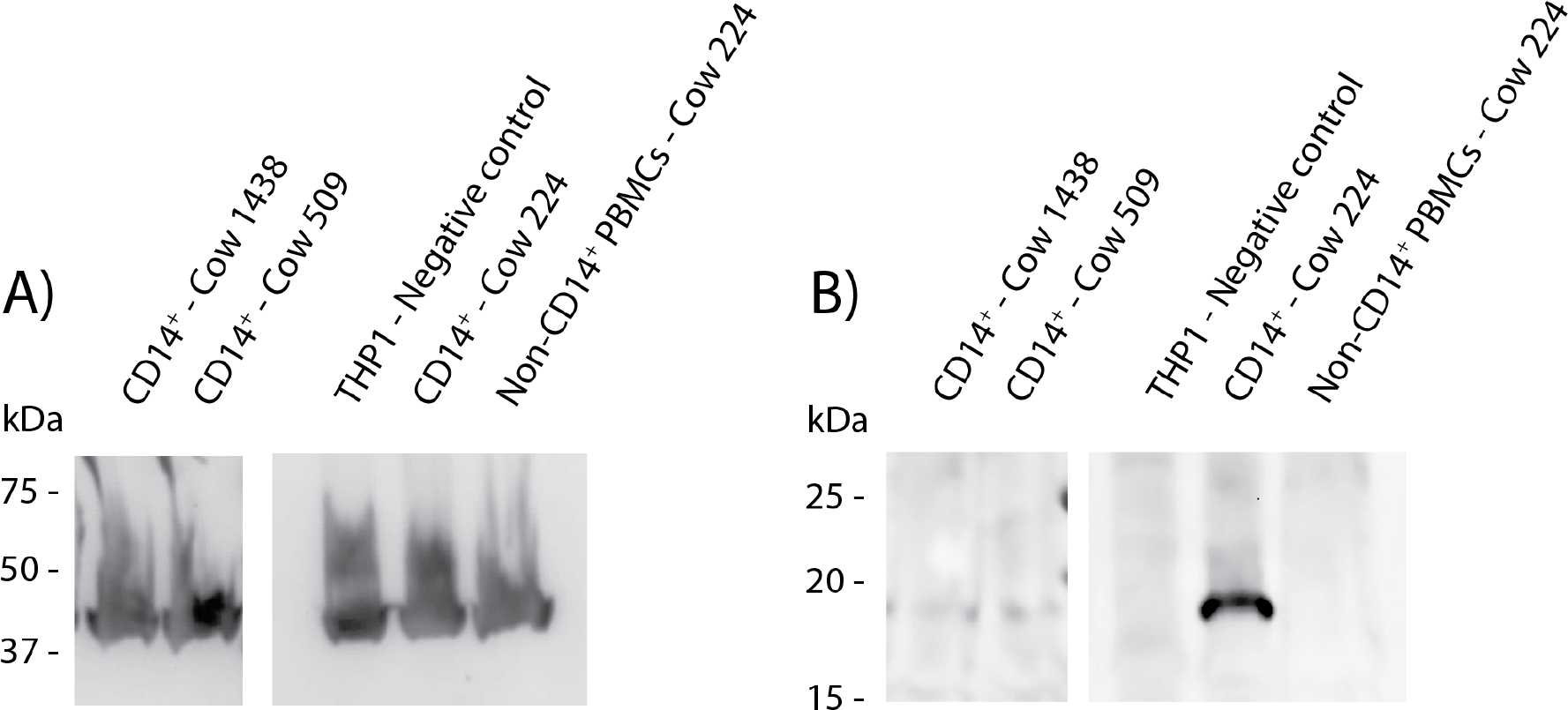
Western Blot analysis to detect full-length Mincle in protein extract from bovine Mo Protein extract was generated from CD14^+^ cells of three cows (1438, 509, 224) as well as THP-1 cells (negative control) and assessed by Western Blot analysis. Protein samples were separated by SDS-PAGE using non-reducing conditions. A) Bovine ß-actin was detected by ECL and functions as a loading control showing approximately equal loading between samples. B) ECL detection of anti-boMincle HuCAL antibody AbD31657 followed by goat anti-human IgG F(ab’)2: HRP was used to detect full-length Mincle (19 kDa) in three samples. All images shown above originate from the same nitrocellulose membrane. Mincle expression levels varied strongly between animals.

### Anti-bovine Mincle HuCAL antibodies react with the full-length bovine Mincle protein expressed either in CHO cells or by bovine leukocyte subsets

The 16 HuCAL antibodies were further assessed by flow cytometry to identify the strongest reacting ones for subsequent experiments. Antibodies were tested on either CHO cells transfected with full-length bovine Mincle in pUNO1 or on bovine PBMCs. An example of the gating strategy for the analysis of Mincle expression on bovine PBMCs is shown in (Suppl. Fig. 5). On CHO cells, positive surface staining was detected on up to 53.32 % of transfected cells (Fig. 4A). No staining was observed on control cells transfected with the empty pUNO1 plasmid (data not shown). All antibodies were subsequently tested on PBMCs (recovered from frozen for 48 h) originally isolated from three different cattle. Staining was only observed in the monocyte population, identified by a high FSC and SSC (Suppl. Fig. 5), with antibodies staining up to 71.8 % of cells (Fig. 4B).

**Fig. 4.**
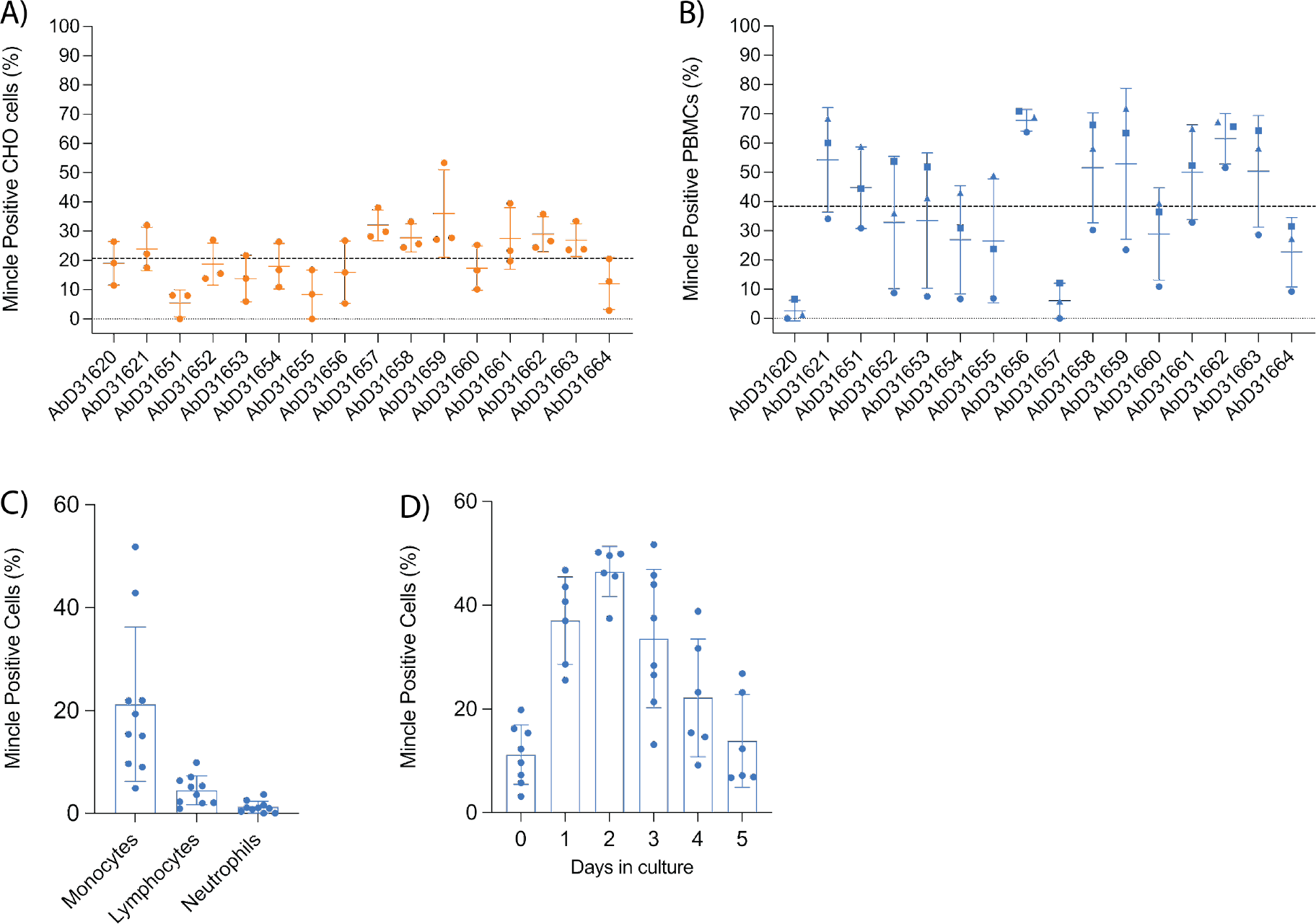
Flow cytometric detection of rboMincle on transfected CHO cells and boMincle on bovine leukocyte subsets and MDMØ A panel of 16 anti-boMincle HuCAL antibodies was analysed by flow cytometry on A) CHO cells transfected with pUNO1 plasmid DNA containing the full-length bovine Mincle gene (CLEC4E) and B) bovine PBMCs isolated from three different cattle. A goat anti-human IgG F(ab’)2: FITC was used as secondary antibody. Dot plots show individual data points for each biological experiment (n = 3). The mean and standard deviation (SD) for each antibody are represented by horizontal bars while the mean positive staining for all data points is represented by a dashed line. Data points representing antibody binding to B) PBMCs from individual cattle are shown as circles (cow 804), squares (cow 1062), and triangles (cow 931). C) Subsequently, anti-boMincle HuCAL antibody AbD31662 was used to detect boMincle on freshly isolated, bovine neutrophils and PBMCs. The scatter dot plot shows the percentage of positive staining for ten individual cattle represented by filled circles. The mean percentage of positive staining for each cell type is represented by bars and the SD by error bars. D) Bovine monocytes, purified from PBMCs by adherence to cell culture plastic, were differentiated into macrophages by addition of M-CSF for one to five days. On each of these five days cells were stained for Mincle using AbD31662 and analysed by flow cytometry. The scatter dot plot shows the percentage of positive staining for six to eight individual cattle at each time point represented by filled circles. The mean percentage of positive staining for each time point is represented by bars and the SD by error bars.

A high degree of variability was observed between the different antibodies when staining both, transfected CHO cells and PBMCs. Antibodies AbD31620 and AbD31657 were found to react well with transfected CHO cells, but not with PBMCs, while the opposite was observed with antibody AbD31651. Additionally, antibody binding to PBMCs varied greatly between cells generated from the three cows (Fig. 4B). Cells from one of the cows showed a much lower overall reactivity with all antibodies compared to cells from the other two cows. Based on the combined data, antibodies AbD31621 and AbD31662 were used for subsequent cell-based experiments. Both antibodies showed above average staining with transfected CHO cells and the PBMCs as well as the lowest SD across the replicated experiments. Furthermore, both antibodies were bovine Mincle-specific, as no reactivity with human PBMCs was observed (Suppl. Fig. 6).

### Bovine monocytes constitutively express Mincle on their surface

We next examined Mincle expression on the surface of freshly isolated, unpermeabilised, primary bovine leukocyte subsets by flow cytometry. Surface expression of Mincle was readily detectable on monocytes from all examined cattle, although expression was found to vary between individual cattle (4.9 to 52.9 %) (Fig. 4C). However, whereas Mincle was constitutively expressed on monocytes, its expression on lymphocytes and neutrophils was either undetectable or only detectable at low levels (<10 % positive staining) (Fig. 4C). In cases where Mincle expression was detected on more that 5 % of the lymphocyte population, further analysis revealed Mincle to be present on both T cells and B cells (Suppl. Fig. 7). Having assessed that Mincle expression seemed to vary between monocytes isolated from different cows, we next assessed whether its expression can be induced by maturation of Mo to MDMØ. To do so, Mo were purified from PBMCs by adherence to cell culture plates and then differentiated into MDMØ by addition of M-CSF. During the differentiation process Mincle continued to be expressed on the surface of the cells, although changes in the level of expression were observed over time (Fig. 4D). Mincle expression increased during the first 2 days of incubation with M-CSF, followed by a decrease over the following 3-4 days to the initial starting levels. This could not be attributed to cell death as analysed by trypan blue exclusion.

For subsequent experiments it was decided that MDMØ cultured with M-CSF for 3-5 days would be used. These cells were also shown to express Mincle and to have characteristics demonstrating a MØ phenotype. In culture, these cells showed the classical “fried egg” morphology, with a spread-out cytoplasm and a more granular appearance compared to freshly isolated Mo and showed increased expression of the MØ markers CD16, CD163, and CD206 compared to Mo, while continuing to express CD14 and MHC class II on their surface (Suppl. Fig. 8).

### Stimulation of bovine MDMØ with Mincle ligands

Since stimulation with LPS and TDB have been shown previously to induce surface expression of Mincle in other mammalian species, we investigated whether this was also the case for bovine MDMØ. Surprisingly, no increase in the surface expression of Mincle was observed following stimulation with LPS, and only a minimal increase (0-5 %) occurred following incubation with TDB (data not shown). Whether this is due to differential receptor expression mechanisms of our monocyte-derived macrophages compared to bone marrow-derived macrophages often used in previous studies (24) remains to be investigated. As no changes in Mincle surface expression were observed as a result of cell stimulation, the role of Mincle in inflammatory responses to these stimuli was investigated. MDMØ from all the cattle tested produced a TNF-α response to LPS and TDB (Fig. 5A). Interestingly, binding of HuCAL antibodies AbD31621 and AbD31662 alone induced a low TNF-α response. Co-incubation of bovine MDMØ with TDB and either of the HuCAL anti-bovine Mincle antibodies had a significant additive effect on TNF-α production (Fig. 5B; p < 0.05), whereas no additional effect was observed when MDMØ were incubated with LPS and either antibody (data not shown).

**Fig. 5.**
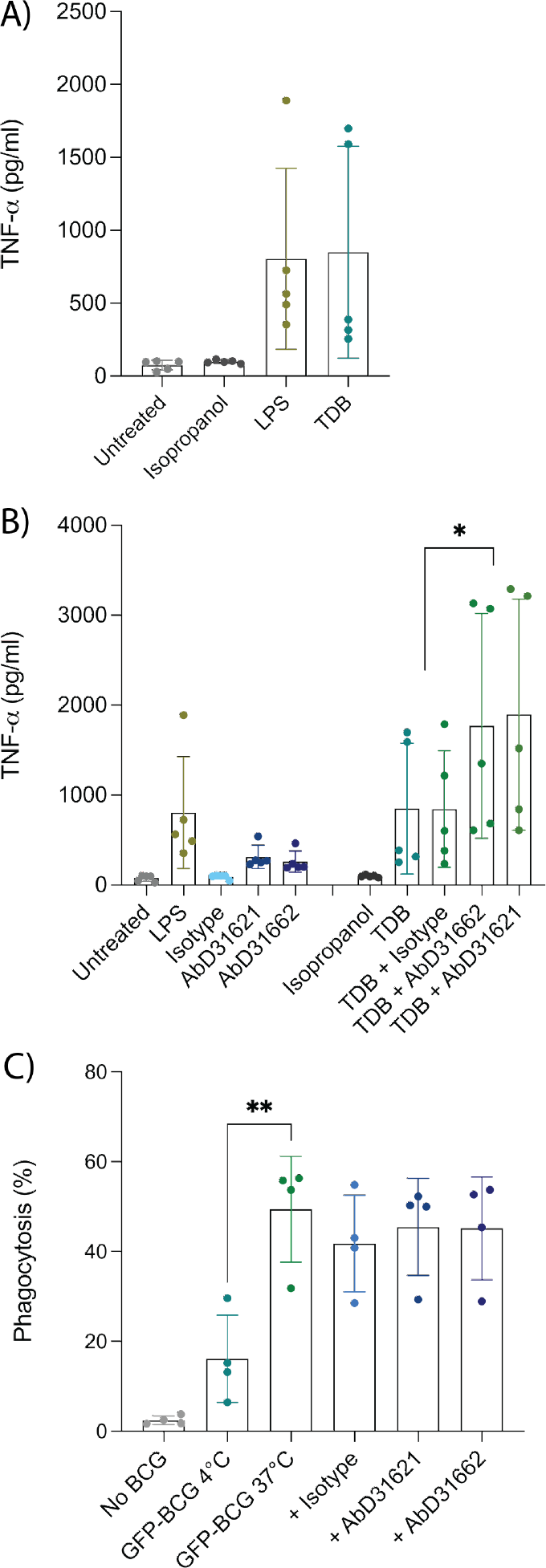
Investigation of the role of Mincle in TNFa production by and phagocytic activity of bovine MDMØ TNF-α secretion by bovine MDMØ was measured by ELISA in cell culture supernatants from cells incubated with (A) LPS or TDB (B) and anti-boMincle antibodies with and without TDB. The scatter dot plots show TNF-α secretion (pg/ml) for cells from five individual cattle represented by filled circles. The mean for each condition is represented by bars and the SD by error bars (p ≤ 0.05 (*)). C) To determine the role of Mincle in the phagocytosis activity of bovine MDMØ, cells were incubated with GFP-labelled BCG (MOI of 5) for 90 min at 37°C, 5 % CO_2_ and analysed by flow cytometry. Cells incubated at 4°C were used as a control. The role of Mincle was investigated by addition of anti-boMincle antibodies. The scatter dot plot shows the percentage of phagocytosis of GFP-BCG by bovine MDMØ from four individual cattle represented by filled circles. The mean for each condition is represented by bars and the SD by error bars (p ≤ 0.01 (**)).

### Phagocytosis of GFP-BCG by MDMØ and blocking with Ab

As Mincle has been described to be involved in the uptake/phagocytosis of *Mycobacterium spp*., we assessed phagocytosis of *M. bovis* BCG stably expressing green fluorescent protein (BCG-GFP) by bovine MDMØ by flow cytometry in the presence of anti-Mincle antibodies (Fig. 5C). Incubation at 4°C was used to assess baseline phagocytosis. Significantly higher amounts of BCG-GFP uptake were observed at 37°C compared to 4°C (p < 0.005). However, none of the antibodies chosen seemed to affect the level of uptake of *M. bovis* BCG-GFP (Fig. 5C).

### Bovine Mincle binds to bovine CD4^+^ T cells via a yet unidentified ligand

Depending on the species investigated, Mincle can react not only with pathogens, but also with DAMPs. To assess whether boMincle has a similar function, we transiently transfected CHO cells with a variety of plasmids, including empty pUNO1, pUNO1-boDC-SIGN, and pUNO1-boMincle. Protein expression was confirmed by flow cytometry (Fig. 4A, Suppl. Fig. 9). The cells were then transferred to chamber slides and left overnight before being incubated with sorted, CSFE-labelled, bovine CD4^+^ T cells and analysed by fluorescence microscopy. The number of fluorescent cells was counted on six randomly selected fields of view for each well of the chamber slide (Fig. 6). The wells containing CHO cells transfected with the empty pUNO1 plasmid and pUNO1-boDC-SIGN served as negative and positive controls, respectively. The effect of receptor blockage was assessed by incubating HuCAL anti-bovine Mincle CRD antibody (AbD31662) with pUNO1-boMincle transfected CHO cells for 1 h prior to incubation with CFSE-labelled T cells.

**Fig. 6.**
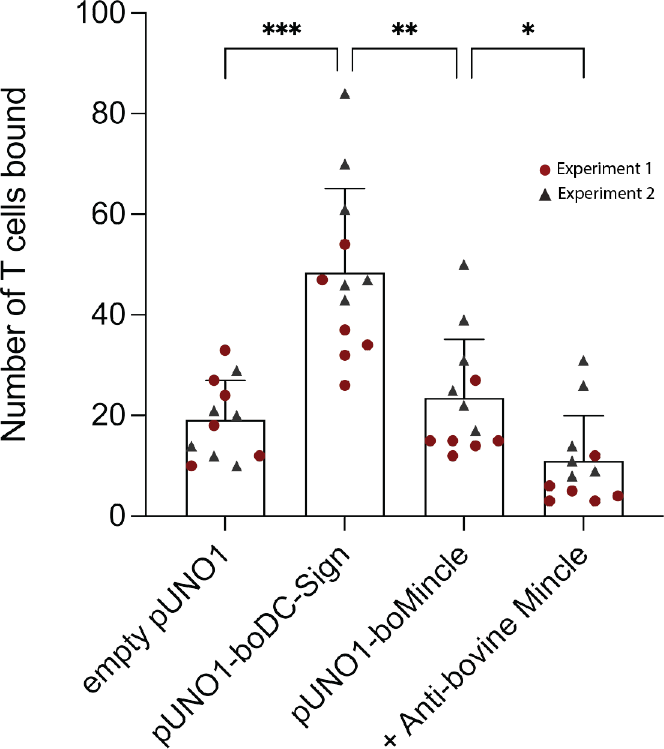
Identification of a potential boMincle ligand expressed on bovine CD4^+^ T cells To identify potential self-derived boMincle ligands, A) CFSE-labelled CD4^+^ T cells were incubated with pUNO1-boMincle-transfected CHO cells. CHO cells were either untreated or pre-incubated with anti-boMincle antibody AbD31662. CHO cells transfected with empty pUNO1 and pUNO1-boDC-SIGN plasmid DNA served as negative and positive control, respectively. The number of CHO cell-bound CD4^+^ T cells was analysed by fluorescence microscopy. The individual results for each of six randomly selected fields of view are represented by red circles (first experiment) and grey triangles (second experiment). The mean for each condition is shown by bars and the SD by error bars (p = 0.0332 (*), p = 0.0021 (**), p = 0.0002 (***)).

As expected, only a few fluorescent T cells bound to CHO cells transfected with the empty pUNO1 plasmid, while CHO cells expressing boDC-SIGN showed significant binding of fluorescently labelled CD4^+^ T cells (p = 0.0002). The number of fluorescently labelled CD4^+^ T cells binding to boMincle-expressing CHO cells was higher compared to CHO cells transfected with the empty plasmid. However, it was also significantly less compared to CHO cells expressing boDC-SIGN (p = 0.0021), despite surface expression of both receptors on CHO cells being similar (Suppl. Fig. 9). Incubation of boMincle-expressing CHO cells with anti-bovine Mincle antibody significantly reduced the binding of fluorescently labelled CD4^+^ T cells (p = 0.0332), thus demonstrating that binding between bovine Mincle and T cells is specific. Repeated attempts to isolate the ligands from CD4^+^ T cells and bovine PBMCs using immunoprecipitation with recombinant boMincle were, however, unsuccessful.

## Discussion

In the present work, we identified the genomic localisation of bovine *CLEC4E*/Mincle within the C-type lectin cluster on bovine chromosome 5 and its homology to other mammalian Mincle receptors. Analysis of the amino acid sequence indicates that the receptor has the structure typical for other C-type lectin receptors (CLRs) while lacking their own intracellular signalling motifs and relying on the immunoreceptor tyrosine-based activation motif (ITAM)-containing adaptor molecule Fc receptor γ-chain (FcRγ). Mincle falls within Group II of the CLR family, and is grouped together with a subset of CLRs termed the Dectin-2 cluster within the telomeric region of the natural killer cell complex (2). Compared to human and murine Mincle, boMincle shows only around 70 % sequence identity, implying that some evolutionary divergence has occurred between species and the ligands that need to be recognised by the host. However, recognition of individual ligands including trehalose and acylated trehalose derivatives is likely similar across these three species (18).

### Expression and regulation of boMincle

We detected boMincle transcripts in all leukocyte subsets, including granulocytes, which is in-line with published literature (8, 25). Furthermore, multiple bands of PCR-amplified boMincle cDNA were observed for sorted leukocyte subsets of some animals, indicating the presence of potential splice variants, an observation also predicted by NCBI. As the primers used for RT-PCR amplified all potential variants, we next assessed whether antibodies raised against the rboMincle CRD would detect the protein by Western Blot and flow cytometry.

In Western Blot analyses, all of the tested anti-rboMincle HuCAL antibodies detected the recombinant bovine Mincle CRD (Suppl. Fig. 4) and a protein of the expected size of full-length Mincle in bovine CD14^+^ Mo (Fig. 3). Interestingly, a particularly strong band of the expected size was observed for protein extract from bovine CD14^+^ Mo from a cow (224) previously infected with the apicomplexan parasite *Theileria (T*.*) parva. T. parva* is a tick-borne hemoprotozoan that infects and transforms bovine lymphocytes (26, 27). Cow 224 had been immunised with a preparation of live *T. parva* sporozoites (“Infection and Treatment Method”) (28), which triggers an antigen-specific, MHC class I-restricted, cytotoxic T cell response among other immune reactions (29, 30). However, *T. parva* infection of cow 224 had been cleared via antiprotozoal drug treatment prior to blood sample collection and PBMC isolation for this work. This finding therefore highlights that the immune status might play an important role in Mincle expression and that such expression can be affected lastingly by past exposure to certain pathogens. It further suggests that Mincle protein expression by CD14^+^ Mo of cattle (1438, 509) unexposed to such pathogens or corresponding stimuli is low as it was only marginally detectable by Western Blotting (Fig. 3).

Flow cytometric evaluation of Mincle expression on the surface of transfected CHO cells with the custom panel of HuCAL antibodies resulted in positive staining of up to half of the cells. In contrast, bovine PBMCs showed antibody staining only on the Mo subset with a considerable variability between individual cows. Notably, these cows belong to the same herd with identical vaccination protocols and pathogen exposure. Two of the antibodies tested reacted only with boMincle and not with huMincle (Suppl. Fig. 6). These two antibodies were used for further evaluation of Mincle expression in the bovine system.

Surface expression of boMincle appeared to depend on the activation status of the respective cells. Consequently, we evaluated Mincle expression levels throughout the differentiation of bovine Mo to MØ. Indeed, Mincle expression increased during the first two days of maturation but decreased over the following three to four days to the initial starting level (Fig. 4D). This couldn’t be attributed to cell death and might have been the result of a lack of appropriate stimuli. Other studies assessing Mincle expression also describe the necessity for cell stimulation to detect Mincle. A study investigating Mincle expression in bovine urothelial cell tumours only detected the receptor in the neoplastic cells by RT-PCR and Western Blot, but not in tissue-derived MØ (31). The authors speculated that expression levels of Mincle were simply too low for detection in non-neoplastic tissue (31). This is in line with our Western Blot analysis showing Mincle mostly in protein extract from CD14^+^ Mo of animal 224 (Fig. 3), a cow previously exposed to the transforming, lymphoproliferative parasite *T. parva*. Similarly, the degree of huMincle expression on neutrophils directly correlated with synergistically-enhanced immune responses. Mincle signalling in response to TDM was dependent on stimulation of the Syk immune pathway (7, 32).

While the recognition of certain ligands is likely conserved across human, murine, and bovine Mincle (18-20, 23), species-specific ligand binding affinities of Mincle have been described as well (24). As the expression, potential downregulation, and ligand binding of boMincle is not yet fully understood, it is possible that boMincle is not expressed under the same conditions as murine or human Mincle and that it possesses a different overall ligand binding spectrum.

### Induction of pro-inflammatory responses by and phagocytosis of boMincle ligands

Ligand binding by Mincle results in activation of the Mincle/Syk/NF-kB circuit, an essential pro-inflammatory pathway which is not unique to Mincle activation. This pathway bridges the innate and adaptive immune system, activates various cell types and primes T and B cell responses. In the present study, we examined the effect of LPS, TDB, and *M. bovis* BCG-GFP on MDMØ responses with regards to boMincle surface expression, TNF-α secretion, and phagocytic activity. Neither of these stimuli led to a detectable increase in boMincle surface expression as analysed by flow cytometry (data not shown). However, all stimuli induced an increased TNF-α secretion by exposed cells (Fig. 5A-B). The increased TNF-α secretion after TDB exposure occurred in the presence of either anti-boMincle antibody, but the same was not observed for LPS. The absence of an enhanced TNF-α production to LPS in the presence of anti-boMincle antibodies can be explained by the fact that LPS signals though the LPS receptor complex, comprising MD2-TLR4-CD14 (33).

### Evidence for endogenous surface-expressed boMincle ligand on CD4^+^ T cells

Mincle is a CLR that recognises foreign and self-derived molecules, including proteins (4), glycoproteins (34, 35) and glycolipids. Indeed, Mincle was shown to recognise both self-derived and non-self-derived lipids (11, 36). In addition, Mincle binds endogenous cholesterol crystals (37), cholesterolsulfate leading to sterile inflammation (38), and beta-glucosylceramide from damaged cells (39). However, recognition of these endogenous ligands clearly depends on the species. Cholesterol crystals were recognised by human yet not by murine Mincle (37). More recently, Mincle was reported to bind a variety of self-derived lipids for Mincle-mediated endocytosis that was independent of clathrin-or caeolin-1-containing vesicles in endothelial cells (40). One of the structurally simplest glycosphingolipids (GSLs), β-glucosylceramide, was shown to bind Mincle on myeloid cells and induce immediate inflammatory responses (39).

Given that the sorted CD4^+^ T cells in our assays were alive, we postulate that boMincle also recognises a ligand on this T cell subset (Fig. 6). This is in line with the glycoprotein/-lipid– lectin hypothesis, which explains how cell wall glycoproteins/-lipids and lectins mediate nanoenvironments (41). Glycosphingolipids are one of the most prominent members of cellular glycoconjugates. They are ubiquitously distributed in lipid microdomains, participate in protein interactions and signalling (42), and were shown to be present in the CD4^+^ T cell membrane (43). However, it is also tempting to assume that some of the CD4^+^ T cells underwent apoptosis at the time of the assay, releasing the Spliceosome-associated protein 130 (SAP130). SAP130 is a subunit of histone deacetylase in the nucleus of live cells but diffuses out of dying or damaged cells into the extracellular milieu (4). SAP130 detection by Mincle triggers a pro-inflammatory response and is correlated with disease severity under some clinical conditions (44). Whether this is also the case for boMincle remains to be investigated.

Overall, the present study describes further characterisation of boMincle, the generation of boMincle-specific antibodies as well as evidence for a self-derived ligand expressed on bovine CD4^+^ T cells.

## Acknowledgements

We would like to thank Profs Kurt Drickamer and Maureen Taylor (Imperial College London) for providing the recombinant bovine Mincle CRD fragments with and without biotinylation as well as their support with drafting of the manuscript.

