## Supplementary material for "Characterisation of the bovine C-type lectin receptor Mincle and potential evidence for an endogenous ligand^1^": Suppl Files

### Supplementary Figures:

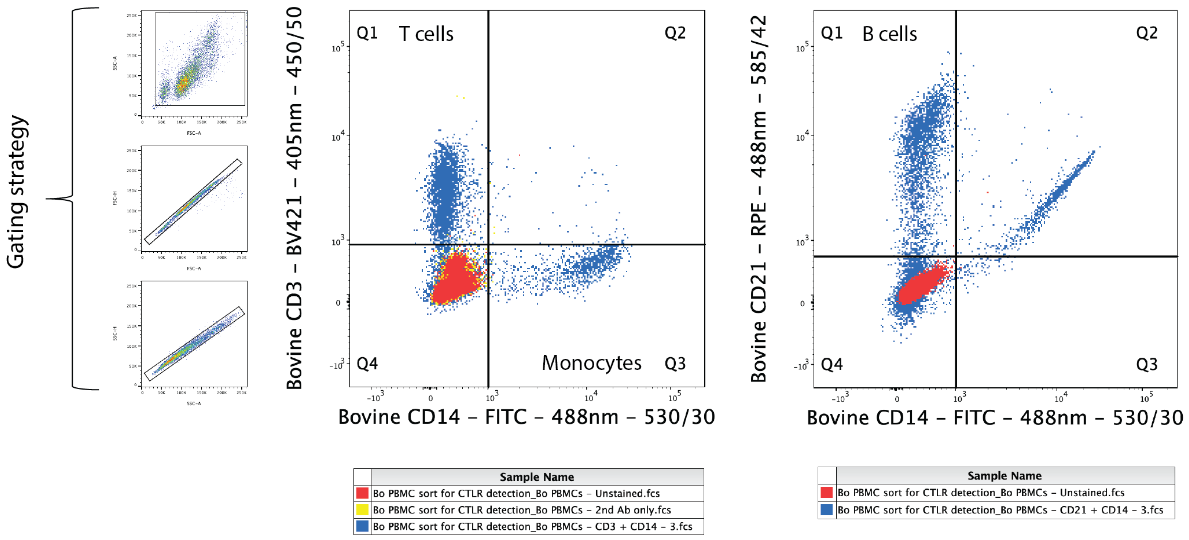


*Supplementary Figure 1: Gating strategy used for FACS of bovine T cells, B cells, and monocytes*

*Bovine PBMCs were either stained for CD3 (T cells) and CD14 (monocytes) or for CD21 (B cells) and CD14 and sorted with a BD FACSAria Fusion cell sorter. Detection of all events was followed by two doublet discriminations (gating strategy) and analysis of the PBMC subsets. Events positive for both RPE and FITC likely occurred due to signal spillover during the analysis and were ignored.*

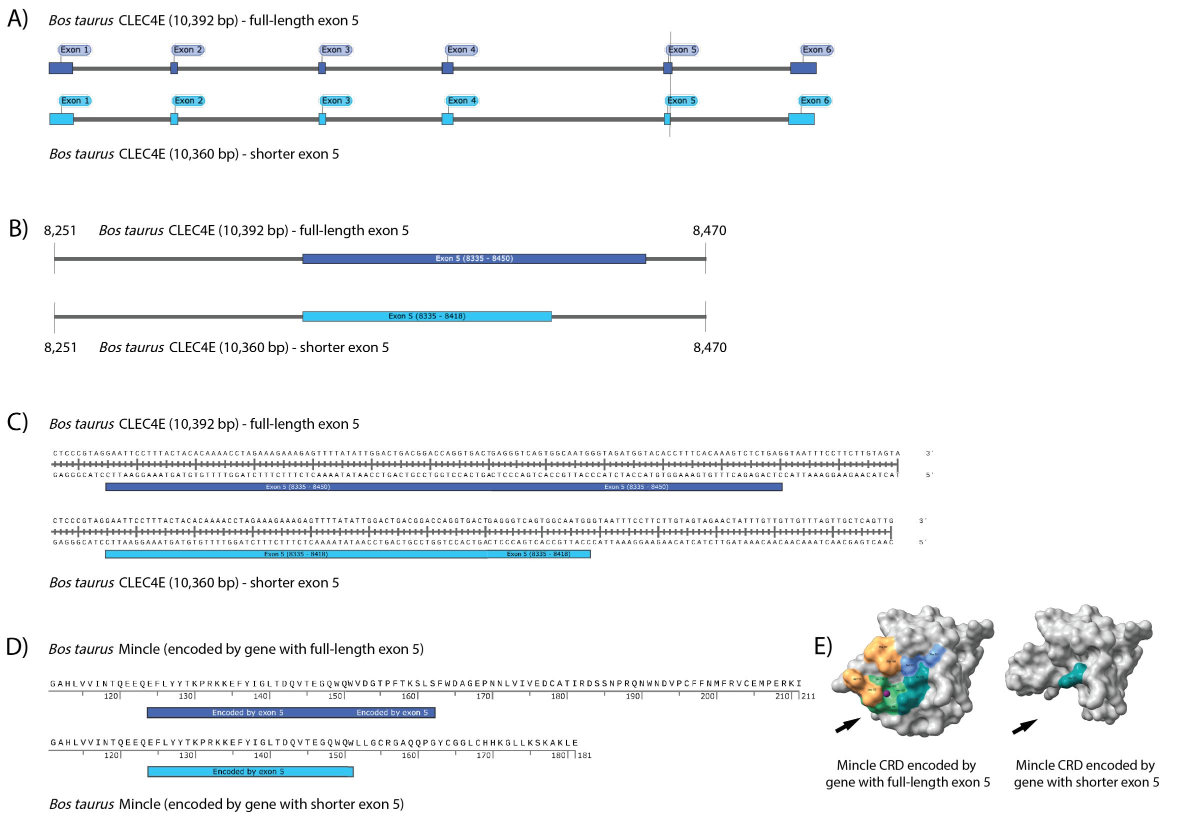


*Supplementary Figure 2: Comparison of CLEC4E gene variants detected in one of the cattle analysed*

*Analysis of polymorphisms in the bovine CLEC4E gene revealed two versions in one of the cattle analysed, A-C) the full-length version and a splice variant missing a 32bp region at the end of exon 5. Potential translation of the splice variant would result in D) a shorter Mincle protein with E) a smaller carbohydrate recognition domain (CRD) that appears to lack the typical carbohydrate binding sites (black arrow) as well as the hydrophobic groove observed in the trehalose-bound CRD of the protein encoded by CLEC4E with full-length exon 5. The colouring of amino acids in both CRD models in E) thereby resembles that shown and described in Figure 2.*

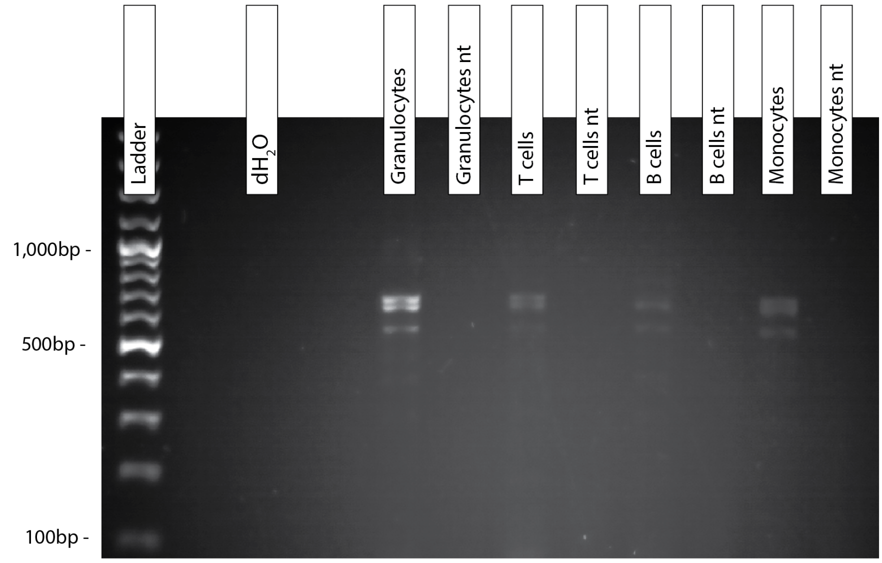


*Supplementary Figure 3: Gel electrophoresis of CLEC4E amplicons after RT-PCR with samples from one of the analysed cattle*

*The transcription pattern of CLEC4E in bovine leukocyte subsets was evaluated by RT-PCR in cDNA samples from freshly isolated, FACS-sorted T cells, B cells, Mo, and granulocytes of up to seven different Holstein Friesian cattle. The image shows the results of the subsequent gel electrophoresis with RT-PCR amplicons of T cells, B cells, monocytes, and granulocytes of one analysed cattle (1132). Samples containing either $dH_2O$ alone or $dH_2O$ instead of RNA and therefore cDNA (nt samples) were included as negative controls. Multiple bands were observed for all four bovine leukocyte subsets which could represent the splice variants of CLEC4E described by NCBI.*

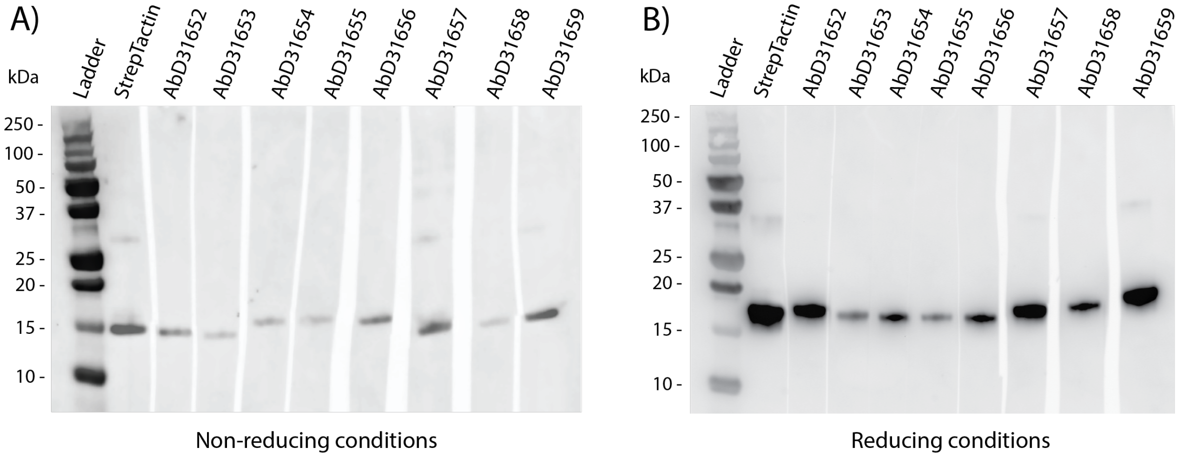


*Supplementary Figure 4: Western Blot analysis to confirm anti-bovine Mincle HuCAL antibody binding of recombinant, bovine Mincle CRD*

*Accurate binding specificities of custom-made, anti-bovine Mincle HuCAL antibodies were assessed by SDS-PAGE and Western Blot analysis using recombinant, bovine Mincle CRD as target. StrepTactin was included as positive control. Staining of nitrocellulose membranes with all tested antibodies resulted in a band of approximately the expected size (15 kDa) both under A) non-reducing and B) reducing conditions.*

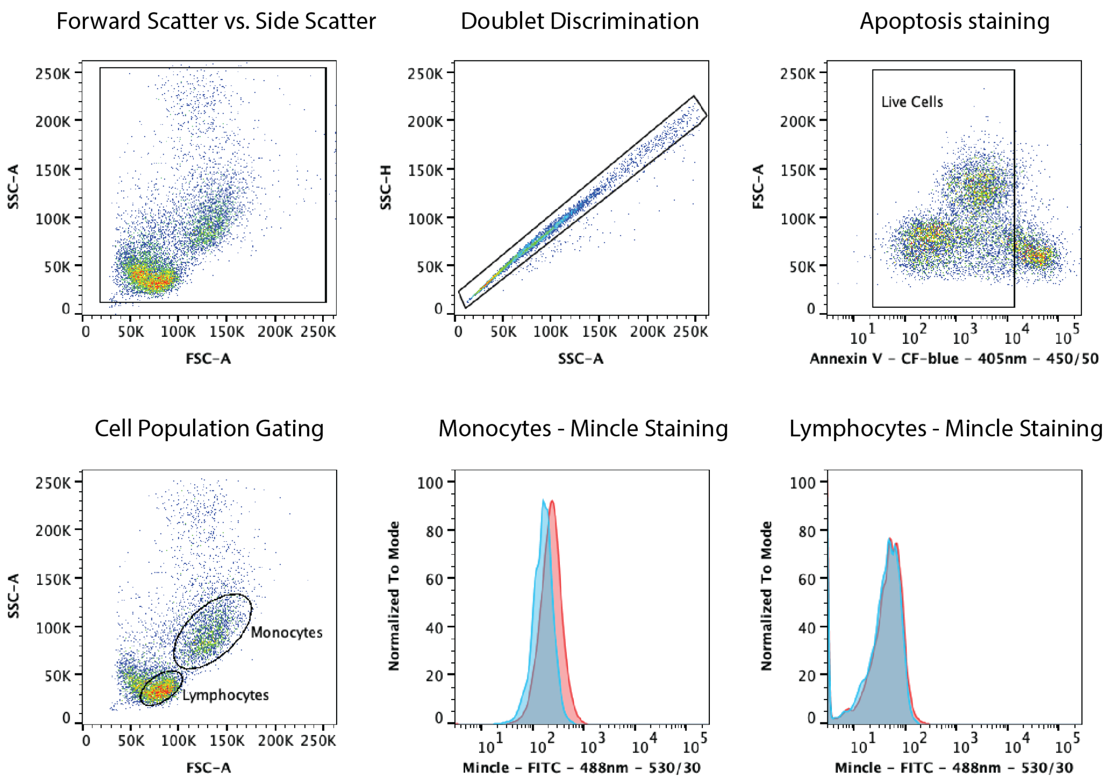


*Supplementary Figure 5: Gating strategy for flow cytometric analysis of Mincle surface expression on bovine PBMCs*

*Bovine PBMCs were labelled indirectly with receptor-specific primary antibodies and a fluorescently labelled secondary antibody and/or with a directly conjugated antibody specific for a cell surface marker. Cell viability was controlled by Annexin V CF-Blue staining. Flow cytometric analysis was carried out on a LSRFortessa X-20. Detection of all events (FSC vs. SSC) was followed by doublet discrimination and selection of live cells. Single, live monocytes and lymphocytes were analysed separately for Mincle surface expression (red histograms) compared to the respective unstained cells (blue histograms).*

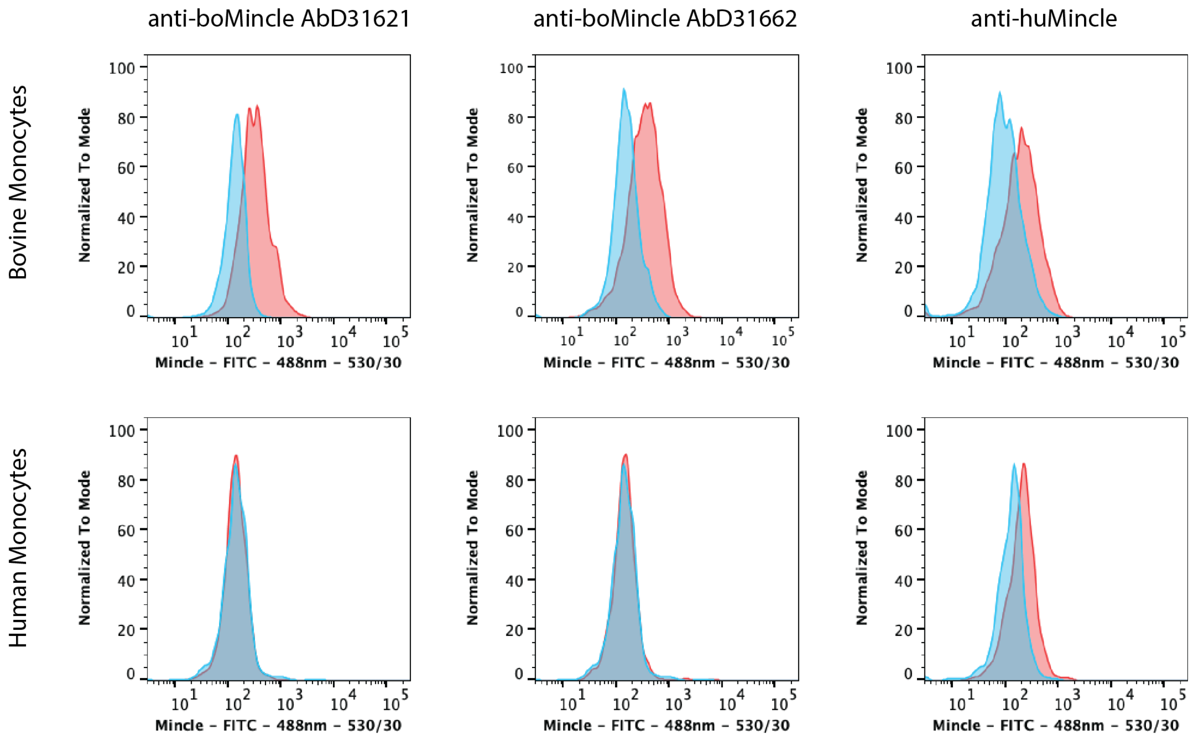


*Supplementary Figure 6: Analysis of cross-reactivity of anti-bovine Mincle HuCAL antibodies*

*Bovine and human monocytes were stained with either anti-bovine Mincle HuCAL antibodies and the goat anti-human IgG F(ab')2: FITC secondary antibody or a FITC-labelled anti-human Mincle antibody for flow cytometric analysis on a LSRFortessa X-20. Single, live monocytes were screened for anti-Mincle antibody staining. The image shows the comparison between surface staining of bovine and human monocytes with anti-bovine Mincle HuCAL antibodies AbD31621 and AbD31662 as well as anti-human Mincle antibody (Invivogen). Stained monocytes (red histograms) were compared to the respective unstained cells (blue histograms). Both bovine-specific antibodies didn't cross-react with human Mincle while the human-specific antibody detected both human and bovine Mincle.*

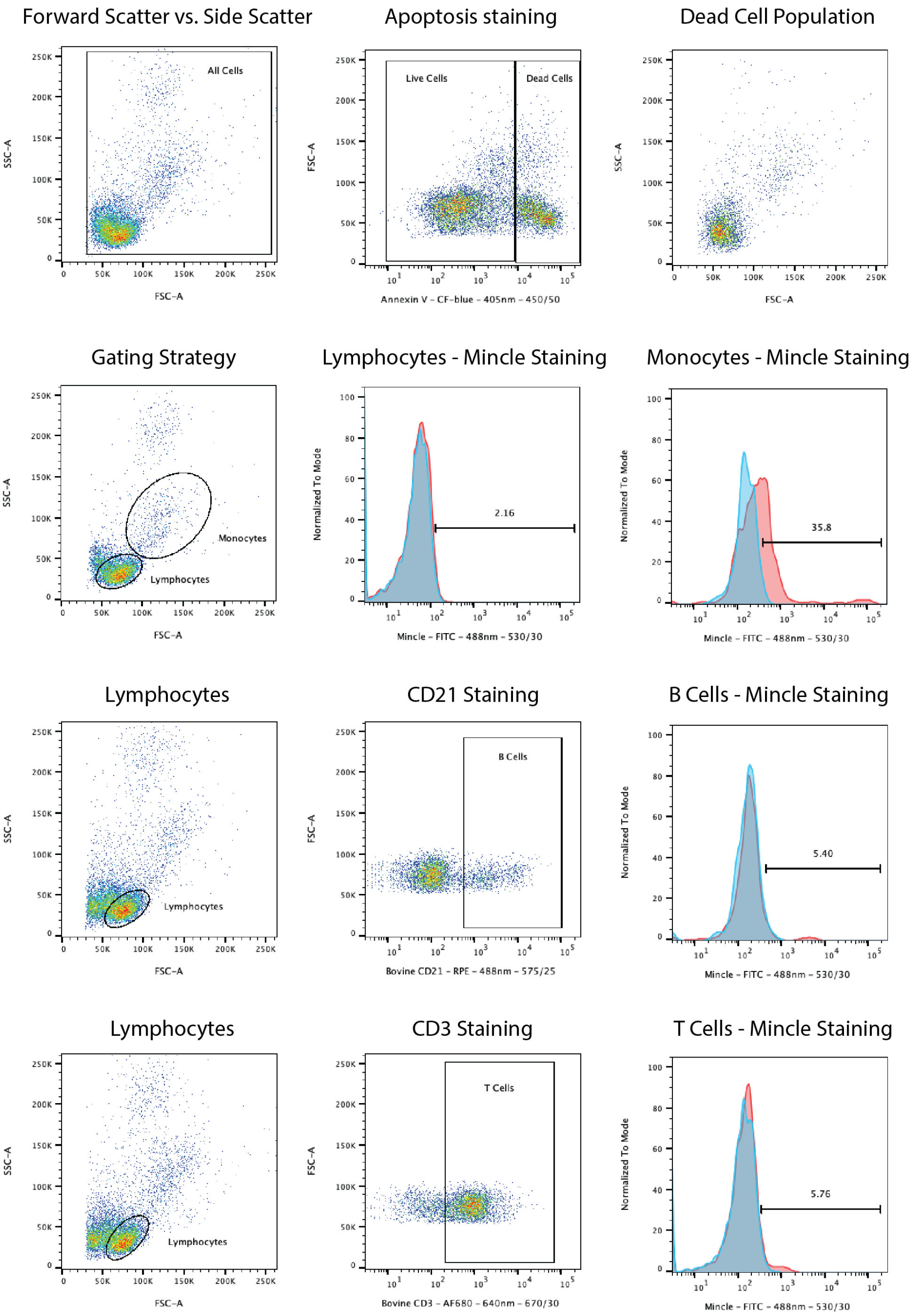


*Supplementary Figure 7: Flow cytometric detection of Mincle on the surface of bovine monocytes, T cells, and B cells*

*Bovine PBMCs were labelled indirectly with receptor-specific primary antibodies and a fluorescently labelled secondary antibody and/or with a directly conjugated antibody specific for a cell surface marker. Cell viability was assessed by Annexin V CF-Blue staining. Flow cytometric analysis was carried out on a LSRFortessa X-20. Monocytes and lymphocyte subsets were analysed separately for surface expression of Mincle. Stained cells (red histograms) were compared to the respective unstained cells (blue histograms). Bovine Mincle was detected on not only monocytes but also more than 5 % of T and B cells.*

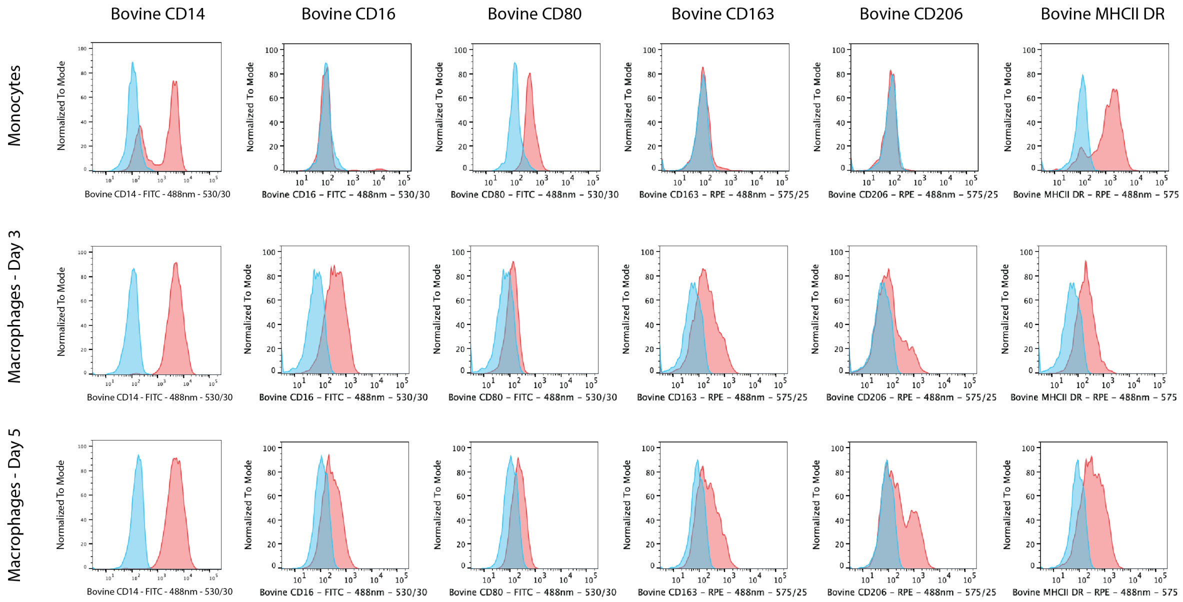


*Supplementary Figure 8: Flow cytometric assessment of bovine Mo differentiation into MØ*

*Bovine Mo were purified from bovine PBMCs by adherence to cell culture plates and differentiated into MDMØ by addition of recombinant M-CSF. The expected maturation of stimulated Mo was assessed by flow cytometric detection of Mo- and MØ-specific surface markers. The image shows surface staining of Mo, day three MØ, and day five MØ (red histograms) for six of these surface markers compared to the respective unstained cells (blue histograms).*

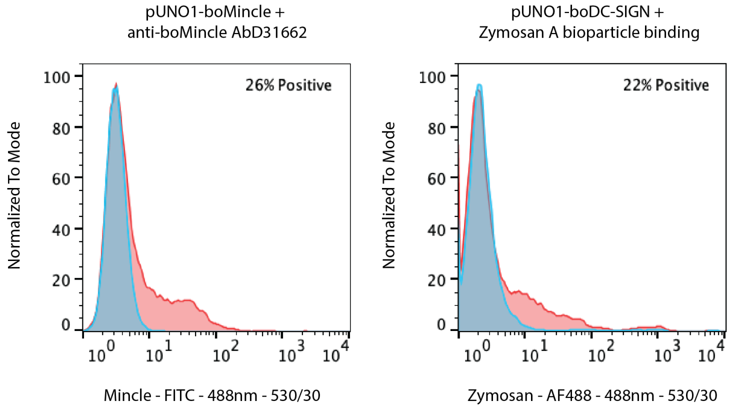


*Supplementary Figure 9: Flow cytometric comparison of boMincle and boDC-SIGN expression on transfected CHO cells*

*CHO cells were transfected with pUNO1-boMincle or pUNO1-boDC-SIGN and analysed by flow cytometry to confirm surface expression of each receptor. Single, live CHO cells stained with either anti-boMincle HuCAL antibody AbD31622 and goat-anti-human IgG F(ab')2: FITC secondary antibody or AF488-labelled Zymosan A bioparticles (red histograms) were compared to the respective unstained cells (blue histograms). Surface expression of both receptors was confirmed and found to be similar (26 % boMincle positive vs. 22 % boDC-SIGN positive CHO cells).*
